# Short-term fasting selectively influences impulsivity in healthy individuals

**DOI:** 10.1101/468751

**Authors:** Maxine Howard, Jonathan P Roiser, Sam Gilbert, Paul W Burgess, Peter Dayan, Lucy Serpell

## Abstract

Previous research has shown that short-term fasting in healthy individuals (HIs) is associated with changes in risky decision-making. The current experiment was designed to examine the influence of short-term fasting in HIs on four types of impulsivity: reflection impulsivity, risky decision-making, delay aversion, and action inhibition. HIs were tested twice, once when fasted for 20 hours, and once when satiated. Participants demonstrated impaired action inhibition when fasted; committing significantly more errors of commission during a food-related Affective Shifting Task. Participants also displayed decreased reflection impulsivity when fasted, opening significantly more boxes during the Information Sampling Task (IST). There were no significant differences in performance between fasted and satiated sessions for risky decision-making or delay aversion. These findings may have implications for understanding eating disorders such as Bulimia Nervosa (BN). Although BN has been characterised as a disorder of poor impulse control, inconsistent findings when comparing individuals with BN and HIs on behavioural measures of impulsivity question this characterisation. Since individuals with BN undergo periods of short-term fasting, the inconsistent findings could be due to differences in the levels of satiation of participants. The current results indicate that fasting can selectively influence performance on the IST, a measure of impulsivity previously studied in BN. However, the results from the IST were contrary to the original hypothesis and should be replicated before specific conclusions can be made.

## Introduction

Impulsivity has been defined as behaviour that can lead to undesirable consequences, is inappropriate to the circumstance, risky, or ill-considered (1). Impulsivity can be categorised into several subtypes, assessed through self-report and behavioural measures [2, 3], and is widely implicated in psychiatric illness (2, 3).

Impulsivity is a multifactorial construct that includes several sub-components, including: reflection impulsivity, action inhibition, delay aversion, and risky decision-making [4, 5].

Reflection impulsivity refers to a reluctance to collect and reflect on information before making a decision, commonly measured using the Matching Familiar Figures Test (MFFT) (4). Participants decide whether figures presented on a screen match one another; the combination of faster and inaccurate responses is associated with higher reflection impulsivity.

Action Inhibition is the failure to inhibit a motor response, and is commonly measured using go/no-go tasks(5). Participants are instructed to respond to target stimuli, and to inhibit responses to distractor stimuli. A greater number of inappropriate responses to the distractor stimuli (commission errors) is usually interpreted as reflecting impaired action inhibition.

Risky decision-making is the tendency to select a larger, but less likely, versus a smaller, but more likely reward and has been measured in a number of different ways, including the Iowa Gambling Task (IGT) (6). The IGT combines different levels of uncertainty, reward, and punishment, hypothesised to mimic real life risky decision-making (7). Participants are asked to chose between four decks of identical looking cards. They are given a hypothetical sum of money and told that some choices will lead to winning money whilst other choices may lead to losing money. However, the decks have predetermined wins so that two decks (disadvantageous choices) are associated with higher rewards, but larger losses, whilst the other two decks (advantageous choices) pay out lower amounts, but rarely lead to a loss. A global score is calculated as the mean number of advantageous choices minus the mean number of disadvantageous choices. Lower scores indicate higher risky decision-making.

Delay aversion has been defined as a preference for smaller rewards sooner vs. larger rewards later [4]. The concept of delay aversion has been captured by tasks such as the Temporal Discounting Task that measures the degree to which an individual is driven by immediate gratification vs. the prospect of a delayed reward (8).

There has been considerable interest in recent years into the impact of periods of fasting on neurocognitive performance (9). Such studies have potential implications for understanding the impact of fasting during diets (particularly those which involve intermittent fasting), or religious fasting such as during Ramadan, as well as potentially for eating disorders. In terms of impulsivity, acute starvation has previously been associated with changes in impulsive behaviour (10). In one study, HIs were more risk seeking after fasting for four hours, compared to when satiated (11). However, other studies find HIs to be more risk averse when fasted compared to satiated (12).

The excessive eating and compensatory behaviours observed in bulimia nervosa (BN) have been understood in terms of problems of impulse control (13). Early research suggested that individuals with BN score higher than healthy individuals (HIs) on self-report measures of impulsivity (13-17). However, the validity of these self-report measures has been questioned (18). More recently, studies have used behavioural tasks to measure different facets of impulsivity in BN. In terms of reflection impulsivity, two studies found that individuals with BN were more impulsive on the MFFT(19). However, another study found no difference between BN and HIs (20). Studies of action inhibition comparing BN and HI have provided mixed results (21). A recent meta-analysis concluded that there was stronger evidence for a deficit of action inhibition for disorder-relevant stimuli (food, and body words/images) in BN compared to standard go/no-go tasks (22). Four studies using the IGT have shown increased risky decision-making for BN when compared to HI (23-26). However, one study found no differences (27). In terms of delay aversion, a recent study showed increased temporal discounting in BN(28).

In summary, recent studies using more objective behavioural measures of impulsivity have shown inconsistent results, suggesting that the clinical stereotype of BN as a disorder of poor impulse control may be an oversimplification (14).

The variable findings in studies examining impulsivity in BN could be accounted for by several factors. Firstly, studies have utilised different tasks, which makes comparisons difficult and limits generalizability (7). Secondly, although researchers have matched HI and BN groups based on Body Mass Index (BMI), a marker of chronic starvation, short-term eating behaviours are not routinely measured. Individuals with BN may engage in acute starvation (short-term fasting) in order to compensate for over-eating (29). As mentioned earlier, acute starvation has previously been associated with changes in impulsive behaviour (9, 10).

Hence, the current study aimed to examine the effect of short-term fasting on performance on well designed and validated tasks measuring four components of impulsivity in HIs, using a within subject, repeated measures design.

In line with the findings that human risk attitudes vary as a function of metabolic state (11, 12), and risk seeking behaviour in animals increases following fasting (10), the primary hypothesis was that (1) short-term fasting would increase risky (i.e. low probability) choices during decision-making. Additionally, the effect of short-term fasting on measures of action inhibition, reflection impulsivity, and delay aversion were explored. It was hypothesised that: (2) short-term fasting would be associated with an increase in commission errors on a task of action inhibition; (3) short-term fasting would decrease the amount of information sampled before making a decision on a task of reflection impulsivity; and (4) short-term fasting would decrease the amount of time individuals are willing to wait to receive a reward during a delay aversion task.

## Method

### Participants

Power calculation for a repeated measures, within subject ANOVA with a small effect size (0.25) and 90% power conducted in G*Power indicated a required sample size of 30. Thirty-three female participants (mean age = 25 years; *SD* = 8.26; range = 18.5-56) were recruited through the University College London (UCL) subject pool. Eligible participants were female, aged 18-50, and had a BMI >18.5. Participants were excluded if: they were currently being treated for any serious medical or psychological condition, including diabetes; they had any history of neurological illness or head injury; or were currently pregnant or breastfeeding. Participants either received course credits or were reimbursed for their time. The research was approved by the University College London Ethics Committee, ref 2337/001. Participants gave written informed consent and a full debrief was provided at the end of the study.

### Procedure

The study used a within-subjects repeated-measures design, assessing behaviour under two conditions: once when participants had fasted for 20 hours; and once when satiated. The mean time between sessions was 7.2 days (*SD* = 1.7, range = 6-16), with each session lasting 90 minutes. During the first session participants underwent the Mini International Neuropsychiatric Interview (MINI), used to assess DSM-IV Axis 1 disorders (30), and completed four behavioural tasks. During the other session participants completed questionnaires and the same behavioural tasks. Task and session order (fasted/satiated) were counterbalanced and randomised. Fasting adherence was assessed using self-reported hunger and blood glucose readings from the distal phalanx area of the index finger using the Freestyle Freedom Lite Blood Glucose Monitoring System, supplied by Abbott Diabetes Care, UK (www.abbottdiabetescare.co.uk). All behavioural tasks were administered on a laptop computer, positioned approximately 60cm from the participant.

Participants were renumerated a the standard university rate for their participation.

## Measures

### Questionnaires

Participants completed: the Beck Depression Inventory (BDI–II, Beck, Steer, Ball, & Ranieri, 1996) a measure of the severity of depressive symptoms; the Eating Disorder Examination Questionnaire-6 (EDEQ-6; Fairburn & Beglin, 1994), to measure ED symptoms; the State-Trait Anxiety Inventory (STAI; Spielberger, Gorsuch, Lushene, Vagg, & Jacobs, 1983), to measure anxiety; and The Impulsive Behaviour Scale (UPPS; Whiteside & Lynam, 2001), to asses impulsivity [39-41]. Additionally, participants filled in a hunger questionnaire that consisted of four Likert scales measuring hunger, desire to eat, the amount of food the participant could eat, and fullness. Participants were also asked to rate from *not at all* to *very much so* how much they were experiencing each of the following: dry mouth, stomach aches, anxiety, dizziness, weakness, nausea, thirst, headache, and stomach growling. A composite score was calculated by adding together the four likert ratings associated with the subjective hunger and the nine ratings of physical side effects. A higher score indicated higher levels of self-reported hunger.

### Experimental Tasks

#### Information Sampling Task (Clark, Robbins, Ersche, & Sahakian, 2006) to measure reflection impulsivity

The Information Sampling Task (IST) measures the degree to which participants sample information before making a decision, whilst placing minimal demands on visual processing and working memory. Participants are shown a 5×5 matrix of 25 grey boxes and are told that each grey box covers one of two possible colours. Participants must decide which colour they think is in the majority, and can click to uncover as many boxes as they wish before deciding. Once opened, boxes remain visible for the remainder of that trial. Correct decisions in the Fixed Win (FW) condition are awarded 100 points, irrespective of number of boxes opened. In the Decreasing Win (DW) condition the number of points to be won decreases by 10 points with every box opened. Therefore in the DW condition participants must tolerate higher uncertainty to win a high number of points as sampling information to a point of high certainty would win few points.

#### Temporal Discounting Task (TDT, Pine, Seymour, Roiser, Bossaert, Friston, Curran, & Dolan, 2009) to measure delay aversion

Temporal discounting is the degree to which individuals discount future rewards, such as deciding whether to spend in the near future or whether to save for the further future, (8). Subjects generally prefer near (spending) to far (saving) rewards, consistent with values being discounted in line with the relevant time delay (temporal discounting). The steeper the discounting, the greater the impulsivity. Participants were asked to choose between two serially presented options of differing magnitude ranging from a monetary value of £1 to £100, and a time delay of one week to one year. The rate at which future rewards are discounted (k) is used as a measure of delay aversion. Participants with a greater discount rate devalue future rewards more quickly. Participants were told that one of the options they chose would be randomly selected and paid for on a pre-paid card with a timed activation date, as used in the original study [23]. However, they were debriefed at the end of the task and no payment was made. The task also contained 20 trials in which one of the choices presented was always larger and available sooner. These ‘catch’ trials were used to determine the subject was paying attention to, and understood the task.

#### Choice x Risk Task (CRT, Rogers, Tunbridge, Bhagwagar, Drevets, Sahakian, & Carter, 2003) to measure risky decision making

The Choice x Risk task is used to investigate three factors thought to affect decision-making: the magnitude of expected gains (reward), the magnitude of expected losses (punishment) and the probabilities of each. On each trial participants were required to choose between two gambles, represented as two bars simultaneously presented on the screen. The amount the bar is filled represents the probability of winning, while wins and losses are displayed numerically at the top and bottom of each bar in green and red text respectively. Participants complete four games, consisting of 20 trials presented in a pseudorandom order. There are eight repetitions of each of 10 trial types, including “gain only” and “loss only” trials. Participants were given 100 points at the beginning of each game and instructed to win as many points as possible. After each trial feedback was given on performance and an updated score was displayed for two seconds.

Standard trial types always contained a control gamble (50/50 chance of winning 10 points) and an experimental gamble. The experimental gamble varies in the probability of winning to be either high or low (75 vs. 25), expected gains are either large or small (80 vs. 20 points) and expected losses either large or small (80 vs. 20 points), producing eight trial types. The other two trial types, ‘gain only’ and ‘loss only’ were used to estimate risk-aversion when choosing between losses, and risk-seeking when choosing between gains. In a ‘gains only’ trial, two options with the same expected value are presented. For example, if participants more frequently choose a 100% chance of a gain of £20 when compared to a 50% chance of gaining £40, they would be exhibiting risk-aversion for gains. Similarly, in a ‘loss only’ trial, two options of equal expected value are presented, such as a 50% chance of a £40 loss, compared to a 100% chance of a £20 loss. If participants are more likely to choose the 50% chance of a £40 loss, they would be exhibiting risk-seeking for losses.

#### Affective Shifting Task (AST, modified from Murphy, Sahakian, Rubinsztein, Michael, Rogers, Robbins, & Paykel, 1999) to measure action inhibition

The AST is a measure of motor inhibitory control. Subjects see pictures from two classes - target and distractor - presented rapidly, one at a time in the centre of the screen. They have to respond to target stimuli by depressing the space bar (go) as quickly as possible, whilst inhibiting responses to distractor stimuli (no-go). The time taken to respond to targets (RTs), failures to respond (omissions), and incorrect responses (commission errors) are recorded, with the latter providing a measure of motor inhibition.

Stimuli were pictures of food (F) or household items (H) taken from an existing database designed for neuropsychological studies of AN (31). Instructions at the beginning of each block indicated which stimulus type to respond to. Each stimulus was presented for 300ms with an inter-trial interval of 900ms. A 500ms/450 Hz tone sounded for each error of commission, but not for omissions. There were 10 blocks (2 practice blocks) with 18 stimuli presented in each block, arranged in either of the following orders: FFHHFFHHFF, HHFFHHFFHH. This order means that four blocks were ‘shift’ blocks, in which participants had to respond to stimuli that were previously distractors, and inhibit responding to previous targets. In the ‘non-shift’ blocks participants had to continue responding to the same targets and inhibiting responses to the same distractors as in the immediately previous block. Note that this was the only one of the included tasks which incorporated food stimuli.

### Statistical Analysis

All statistical analyses were performed using SPSS 21 (IBM SPSS, 2010, Chicago, IL, USA). Two tailed statistical significance was determined as p < 0.05. Descriptive statistics (mean and standard deviations) were calculated for all demographic and questionnaire scores.

#### Information Sampling Task

To investigate the effect of fasting on the amount of information sampled during the IST, the dependent variable, average number of boxes opened before making a decision, was entered into a multivariate analysis. A mixed model ANOVA with the within-subject factors of Session (fasted, satiated), Condition (Fixed Win, Decreasing Win) and the between-subject factor of Order (FW-DW, DW-FW) was conducted separately on the primary outcome of average boxes opened, and the secondary outcome of errors. Any significant interactions were then explored with Bonferroni corrections applied.

#### Temporal Discounting Task

Impulsive choice was calculated as the number of sooner options chosen by each participant, for each trial, separately for the fasted and satiated sessions. A pairwise comparison was used to examine any differences across fasted and satiated sessions.

Maximum likelihood estimation was used in order to calculate the maximum likelihood parameters for the discount rate (*k*), and utility concavity (*r*). For each of the 220 choices for each participant a Bernoulli likelihood (based on the sigmoid of the difference in discounted value) was calculated for the chosen option). Likelihood maximization proceeded via optimization functions in Matlab (The MathWorks Inc., Natick, MA, United States). See Pine and colleagues’ (2009) for further information and methods. Pairwise comparisons were run to examine any differences in the discount rate (*k*), or utility concavity (*r*), between fasted and satiated sessions.

#### Choice x Risk Task

To examine the effect of fasting on risky decision-making, multivariate analysis was conducted on the number of times participants choose the experimental, over the control, gamble (proportionate choice) and the mean deliberation times associated with these choices. The proportionate choices were arcsine transformed prior to statistical analysis in line with Rogers, (32). However, all values presented in tables are untransformed scores, for ease of interpretation.

The proportionate choices were analysed using a within subjects repeated measures 2 x 2 x 2 x 2 ANOVA with the factors of session (fast vs. satiated), probability (high vs. low), expected gains (large vs. small), and expected losses (large vs. small). This ANOVA was then repeated with mean deliberation times (ms) as the dependent variable.

The ‘gains only’ and ‘losses only’ trials were analysed using a within subjects repeated measures 2 x 2 ANOVA with session (fast vs. satiated), and trial type (‘gains only’ vs. ‘losses only’). Analysis was conducted on both proportion and deliberation times separately.

#### Affective Shifting Task

To determine the effect of fasting on performance during the AST, multivariate analyses were conducted separately on reaction times (ms), errors of commission, and errors of omission using a 2 x 2 x 2 repeated measures ANOVA with Stimuli (food, household); Condition (shift, non-shift); and Session (fast, satiated) entered as within-subject factors. Any significant interactions were then explored and the Bonferroni correction was applied.

## Results

Demographic characteristics and questionnaire scores are displayed in Table 1.

**Table 1.** Means and standard deviations for demographic variables and trait measures (*n* = 33),

| | | Mean ± SD | |
| --- | --- | --- | --- |
|  | *Demographic Variables* |  |  |
| | Age (years) | 25.42 ± 8.26 | |
| | Body Mass Index (BMI) | 21.65 ± 3.22 | |
|  | *Trait Measures* |  |  |
| UPPS-P = the | UPPS-P | 85.18 ± 11.55 | Impulsive |
| Behaviour Scale; | BDI | 5.15 ± 4.87 | BDI = Beck |
| Depression | EDE-Q | 7.97 ± 6.45 | Inventory; EDE-Q |
| = Eating Disorder | Trait-STAI | 39.30 ± 9.96 | Examination |
| Questionnaire-6; |  |  | STAI = State-Trait |
| Anxiety Inventory |  |  |  |

### Physiological Analysis

#### Blood Glucose

Pairwise comparisons revealed a significant difference for blood glucose levels between fasting and satiated sessions *t*(32)= −5.07, *p*<0.001. Blood glucose levels in the fasted session (*M*=4.06, *SD*=0.51) were lower than in the satiated session (*M*=4.90, *SD*=871).

### Information Sampling Task

Accuracy scores for identifying the correct box colour were examined and any participants with accuracy scores lower than 60% were excluded from further analysis, in line with the original study [31].

#### Boxes Opened

There was a significant main effect of Session [F(1,28)=9.72, *p*=0.004], a significant main effect of Condition [*F*(1,28)=76.16, *p*<0.001] and a significant Session x Condition interaction [*F*(1,28)=4.49, *p*<0.05]. There was no significant effect of Condition Order for the fasting [*F*(1,28)=0.008, *p*=0.928] or satiated Session [*F*(1,28)=0.284, *p*=0.599]. Pairwise comparisons revealed that participants opened significantly fewer boxes in the DW condition, compared to FW for both fasting *t*(30)=7.86, *p*<0.001 and satiated *t*(30=6.78, *p*<0.001) sessions, see Table 2 for mean scores.

**Table 2.**
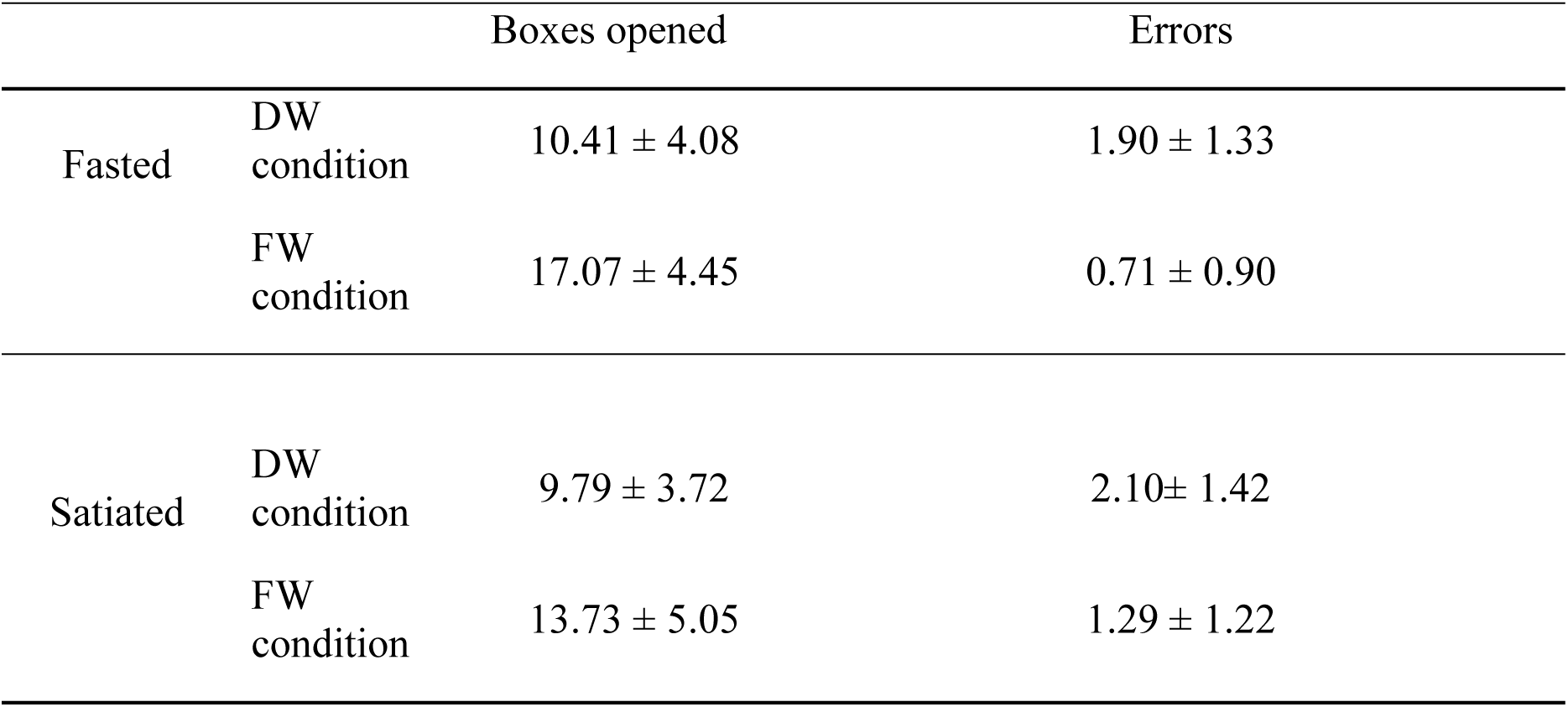
Mean difference and standard deviation (±) scores across fasted and satiated sessions

Post-hoc analysis revealed a significant difference between sessions in the FW condition *t*(30)=3.81, *p*=0.001 but not the DW condition *t*(30)=1.41, *p*=0.168. During the FW condition participants opened more boxes before making a decision, when fasted (*M*= 17.07, *SD*=4.45) compared to when satiated (*M*=13.73, *SD*=5.05).

#### Errors

Analysis of error data using a mixed model ANOVA showed a significant main effect of Session [*F*(1,28)=5.75, *p*<0.05], and a significant main effect of Condition[*F*(1,28)=22.21, *p*<0.001]. The Session x Condition interaction was not significant [*F*(1,28)=0.744, *p*=0.396]. Participants made a higher number of errors during the satiated session, and more errors during the DW condition, see Table 2 for mean scores and standard deviations.

### Temporal Discounting Task

Two participants scored under 80% on the catch trials across both sessions and were therefore excluded from further analysis. All other participants had high accuracy (mean = 19.15) on the catch trials (out of a possible 20). Participants varied on the number of trials in which the sooner option was chosen, ranging from 2 to 184, out of a possible 200 trials. The model of best fit from Pine et al., (2009) showed that participants discounted the value of future rewards (mean fasted *k* = 0.06, SD = 0.68; mean satiated *k* = 0.07, SD = 0.066) and demonstrated a concave utility function (mean fasted *r* = 0.0213, SD = 0.03609; mean satiated *r* = 0.0140, SD = 0.02830). However, the discount rate *t*(30)= −0.521, *p*=0.606 and concave utility *t*(30)= 1.438, *p*=0.161 were not significantly different between fasted and satiated sessions. The impulsive choices made did not differ across session *t*(30)= −0.327, *p*=0.746.

### Choice x Risk Task

Data from three participants were missing for the Choice x Risk Task due to a recording error; therefore 30 participants were included in the following analyses.

### Probability, Wins, and Losses

#### Proportionate Choice

There was no main effect of Session (fasted, satiated) on the proportion of times that participants chose the ‘experimental’ gamble over the ‘control’ gamble [*F*(1,29)=0.22, *p*=0.643]. However, participants chose the ‘experimental’ gamble significantly more often when the probability of winning was high compared to when it was low, [*F*(1,29)=204.73, *p*<0.001], significantly less often when the expected losses were large compared to small [*F*(1,29)=32.95, *p*<0.001], and significantly more often when the expected gains were large compared to when they were small [*F*(1,29)=28.30, *p*<0.001]. However, there was no significant interaction that involved Session (fasted vs. satiated).

#### Deliberation Times

There was no main effect of Session [*F*(1,29)=1.41, *p*=0.26], Probability [*F*(1,29)=1.90, *p*=0.18], or Expected Gains [*F*(1,29)=0.34, *p*=0.57], but a significant main effect of Expected Losses [*F*(1,29)=8.72, *p*<0.01]. Participants took longer to choose when then ‘experimental’ gamble was associated with large expected losses compared to small losses. Means and standard deviations are presented in Table 3. There was no significant interaction that involved Session (fasted vs. satiated).

**Table 3.** Proportion of choices of the ‘experimental’ over the ‘control’ gamble for the probability of winning, expected losses and gains across fasted and satiated sessions

| Group | Probability of winning on the ‘experimental’ gamble |  | Levels of expected losses on ‘experimental’ gamble |  | Levels of expected gains on ‘experimental’ gamble |  |
| --- | --- | --- | --- | --- | --- | --- |
|  | High | Low | Large | Small | Large | Small |
| Fasted | $0.77 \pm 0.33$ | $0.18 \pm 0.18$ | $0.45 \pm 0.25$ | $0.62 \pm 0.21$ | $0.59 \pm 0.23$ | $0.48 \pm 0.20$ |
| Satiated | $0.78 \pm 0.30$ | $0.14 \pm 0.13$ | $0.46 \pm 0.22$ | $0.61 \pm 0.18$ | $0.58 \pm 0.20$ | $0.48 \pm 0.18$ |

**Table 4.**
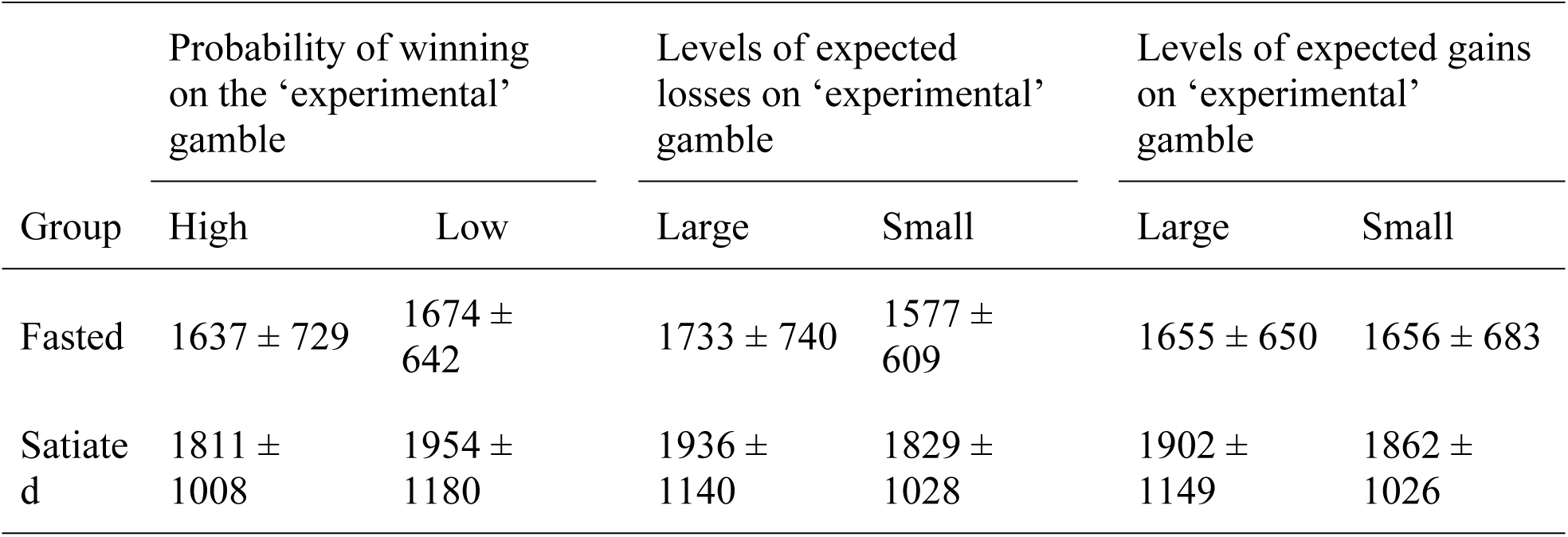
Mean deliberation times (ms) and standard deviation scores for probability of winning, expected losses and gains across fasted and satiated sessions

SHORT-TERM FASTING AND IMPULSIVITY
| Group | Probability of winning on the 'experimental' gamble |  | Levels of expected losses on 'experimental' gamble |  | Levels of expected gains on 'experimental' gamble |  |
| --- | --- | --- | --- | --- | --- | --- |
|  | High | Low | Large | Small | Large | Small |
| Fasted | 1637 ± 729 | 1674 ± 642 | 1733 ± 740 | 1577 ± 609 | 1655 ± 650 | 1656 ± 683 |
| Satiated | 1811 ± 1008 | 1954 ± 1180 | 1936 ± 1140 | 1829 ± 1028 | 1902 ± 1149 | 1862 ± 1026 |

### ‘Gains Only’ vs. ‘Losses Only’ Trials

#### Proportionate Choice

Participants chose the guaranteed options significantly more often on the ‘gains only’ trials compared to the ‘losses only’ trials [*F*(1,29)=83.07, *p*<0.001]. Overall choice was unaffected by Session [*F*(1,29)=0.41, *p*=0.53] and the interaction between session and trial type was non-significant [*F*(1,29)=0.85, *p*=0.77].

#### Deliberation Times

Participants were significantly faster to choose on the ‘gains only’ trials compared to the ‘losses only’ trials [*F*(1,29)=12.34, *p*=0.001]. Reaction times were unaffected by Session session [*F*(1,29)=1.11, p=0.30] and the interaction between session and trial type was non-significant [*F*(1,29)=0.314, p=0.58].

### Affective Shifting Task

#### Reaction Time

There was a significant main effect of Stimuli [*F*(1,32)= 15.26, *p* < 0.001], and Condition *F*(1,32)= 5.38, *p* < 0.05, but no significant effect of Session [*F*(1,32)=0.25, *p* = 0.617]. There was no significant interaction between: Session and Condition [*F*(1,32)= 1.76, *p* = 0.194]; Session and Stimuli (*F*(1,32)= 1.34, *p* = 0.26); Condition and Stimuli [*F*(1,32)= 0.48, *p* = 0.49]; or between Session, Condition and Stimuli [*F*(1,32)= 0.08, *p* = 0.78].

Overall, reaction times (RTs) for food stimuli were shorter (*M*=462.65, *SD*=57.89) than for household items (*M*=482.02, *SD*=56.70). Non-shift trials also had shorter RTs (*M*=468.44, *SD*=57.55), compared to shift trials (*M*=476.24, *SD*=57.04).

#### Errors of Commission

There was a significant main effect of Session [*F*(1,32)= 5.39, *p* < 0.05] but not of Stimuli [*F*(1,32)= 0.15, *p* = 0.69]. There was also a significant main effect of Condition [*F*(1,32)= 43.5, *p* < 0.001]. The interaction between Session and Stimuli was not significant [*F*(1,32)= 2.88, *p* = 0.10], nor was the interaction between Session and Condition [*F*(1,32)= 0.27, *p* = 0.610], or Stimuli by Condition [*F*(1,32)= 0.16, *p* = 0.695]. However there was a significant interaction between Session, Stimuli, and Condition [*F*(1,32)= 4.82, *p* = *p* < 0.05].

More commission errors were made during the fasted session (*M*=1.55, *SD*=0.89), than the satiated session, (*M*=1.19, *SD*=0.82). Participants also made a higher number of commission errors for shift (*M*=1.41, *SD*=1.02), compared to non-shift conditions (*M*=0.14 *SD*=0.81).

Bonferroni post hoc comparisons to explore the Session by Stimuli by Condition interaction showed that there was no difference in the number of commission errors made towards household items between fasted and satiated sessions, for either shift (*p*= 0.33) or non-shift (*p*=0.23) blocks. There was also no difference in commission errors towards food stimuli for fasted or satiated sessions during the non-shift block (*p* = 0.44). However, there was a significant difference in the number of commission errors in response to food stimuli during the shift blocks (*p* < 0.05). There was a higher number of commission errors in response to food stimulus during fasted (*M*=2.39, *SD*=2.21) compared to satiated sessions (*M*=1.36, *SD*=1.48), see Figure 1.

**Figure 1.**
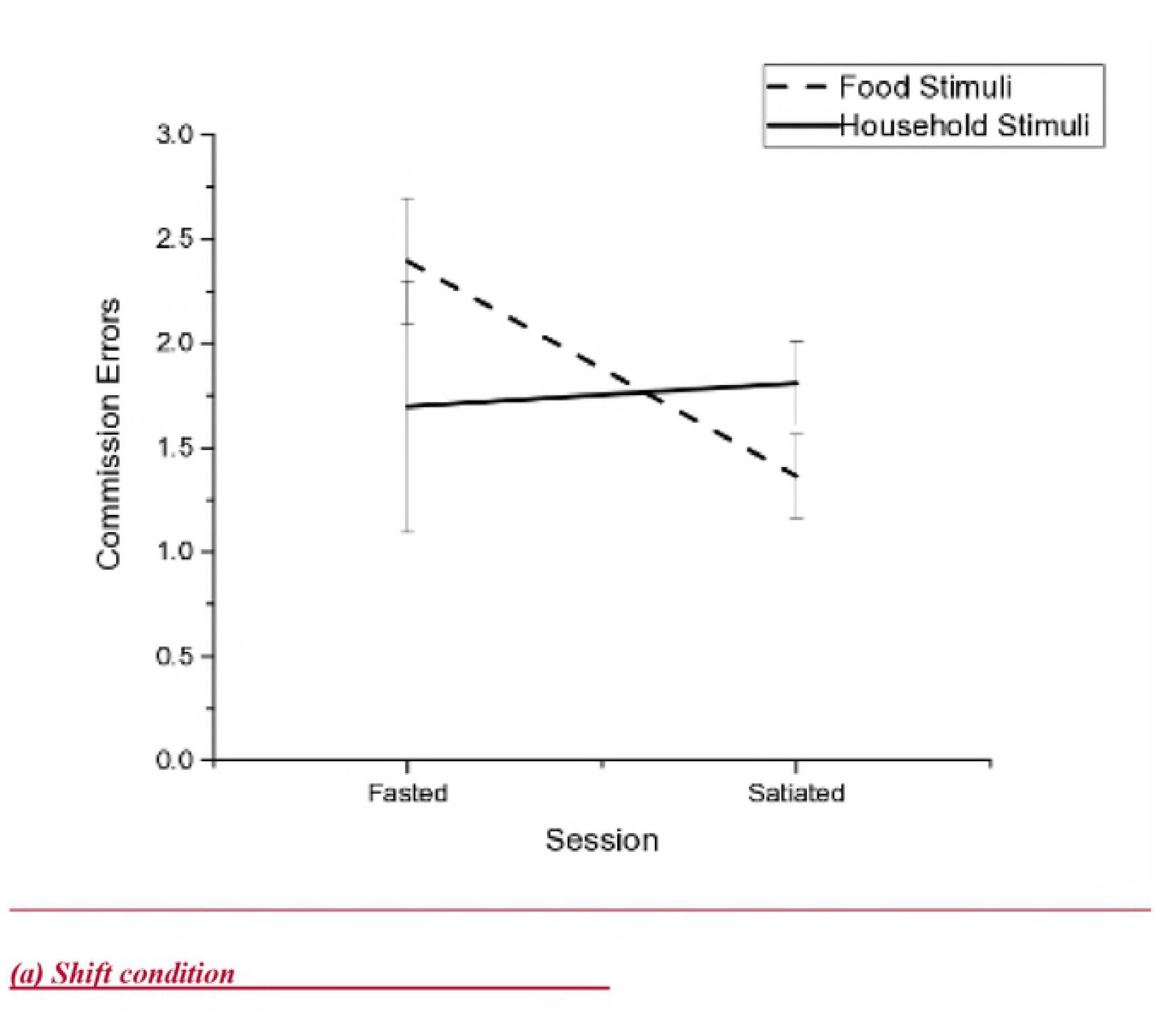
Mean number of commission errors made during the Affective Shifting Task for food and household stimuli across fasted and satiated sessions.

#### Errors of Omission

There was no main effect of Session *F*(1,32)=0.62, *p* = 0.44 or Stimuli *F*(1,32)=0.005, *p* = 0.95. However, there was a significant main effect of Condition *F*(1,32)= 6.17, *p* < 0.05. The interaction between Session and Stimuli was not significant, *F*(1,32) = 0.88, *p* = 0.36, nor was the interaction between Stimuli and Condition *F*(1,32)= 0.25, *p* =0.62, nor the interaction between Session, Stimuli, and Condition *F*(1,32)= 0.42, *p* = 0.517. There was a significant interaction between Session and Condition *F*(1,32)= 7.52, *p* < 0.05. Participants made a higher number of errors of omission during shift blocks (*M*=1.06, *SD*=0.90), compared to non-shift blocks (*M*=0.77, *SD*=0.87). The Session by Condition interaction was explored using Bonferroni adjusted comparisons and revealed that participants made more errors of omission during shift blocks when satiated (*p* < 0.05). However, there was no difference in omission errors between shift and non-shift blocks when fasted (*p* = 0.44).

### Relationship between Self report and Behavioural Measures

Change scores between satiated and fasted sessions were calculated for the commission errors made during the AST, and for the number of boxes opened during the FW condition of the IST. Change scores for the state self report measures were also calculated (state anxiety, blood glucose, and hunger). Correlations between these variables were then calculated. However there was no significant correlation between the self report measures and difference scores for the IST and AST. See Table 5.

**Table 5.**
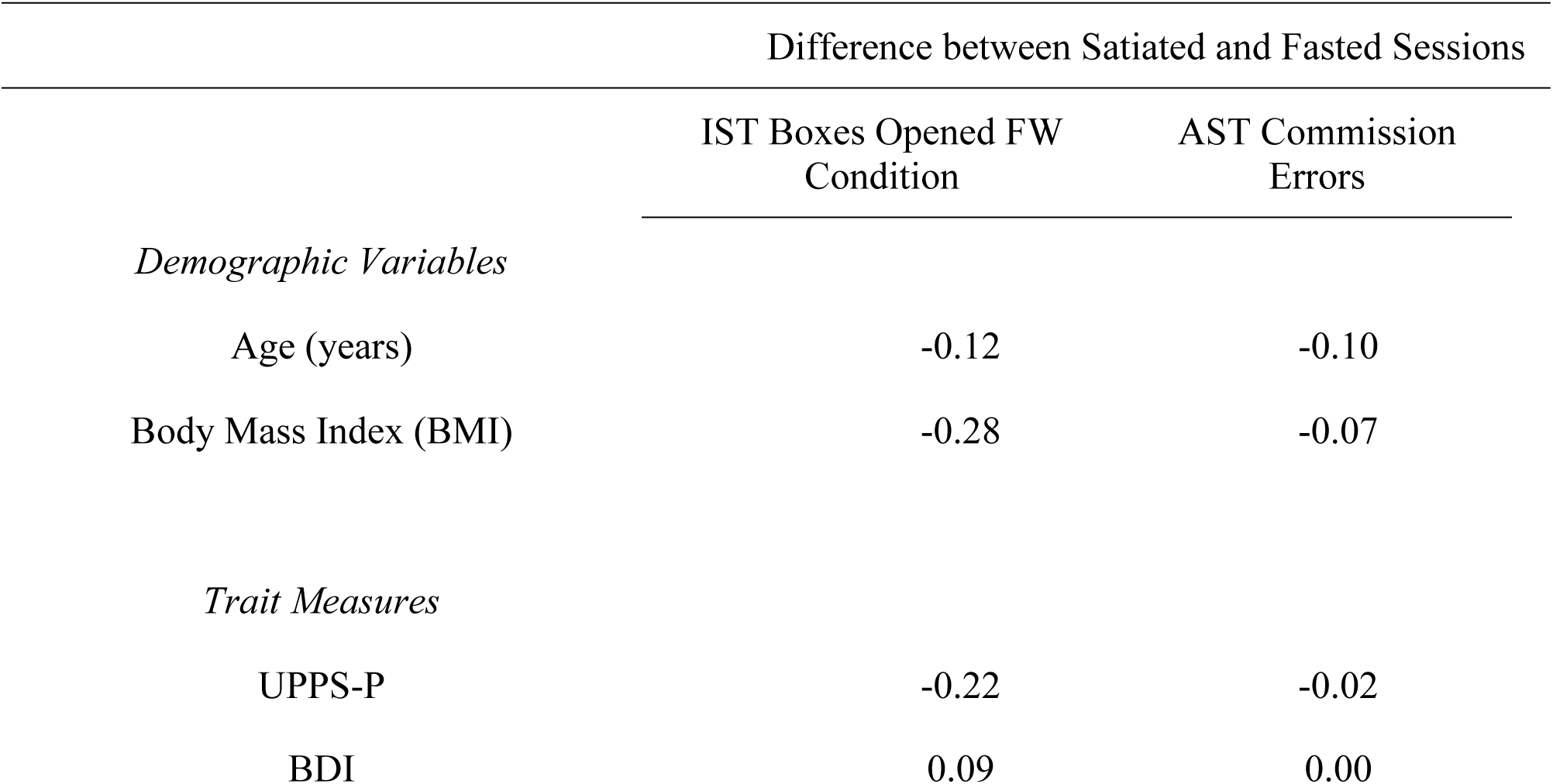
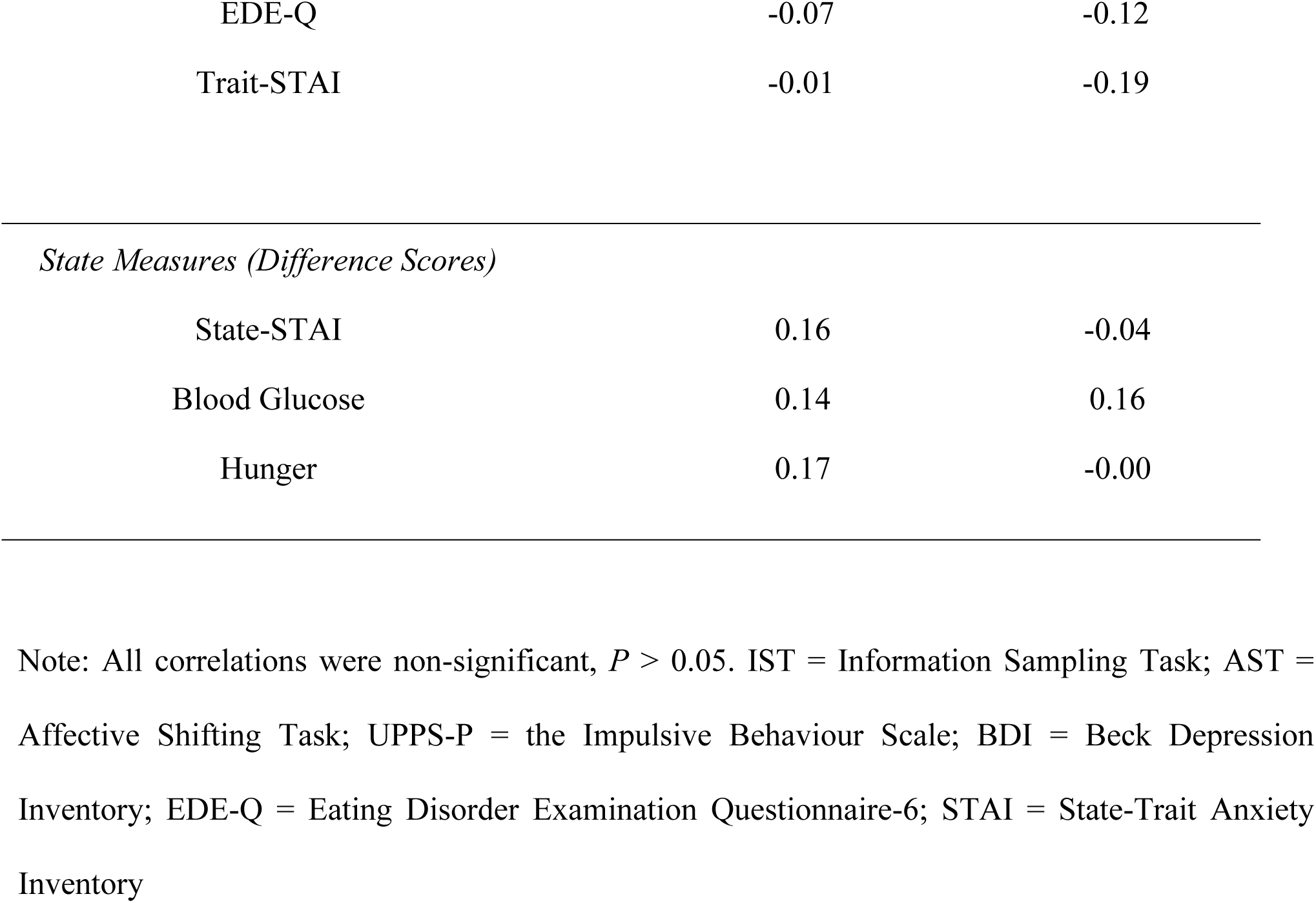
Pearson correlations between the IST and AST difference scores (satiated minus fasted) and state changes in Anxiety, Blood Glucose and Hunger.

|  | Difference between Satiated and Fasted Sessions |  |
| --- | --- | --- |
|  | IST Boxes Opened FW<br>Condition | AST Commission<br>Errors |
| *Demographic Variables* |  |  |
| Age (years) | -0.12 | -0.10 |
| Body Mass Index (BMI) | -0.28 | -0.07 |
| *Trait Measures* |  |  |
| UPPS-P | -0.22 | -0.02 |
| BDI | 0.09 | 0.00 |
| EDE-Q | -0.07 | -0.12 |
| Trait-STAI | -0.01 | -0.19 |
| *State Measures (Difference Scores)* |  |  |
| State-STAI | 0.16 | -0.04 |
| Blood Glucose | 0.14 | 0.16 |
| Hunger | 0.17 | -0.00 |

Note: All correlations were non-significant, $P > 0.05$ . IST = Information Sampling Task; AST = Affective Shifting Task; UPPS-P = the Impulsive Behaviour Scale; BDI = Beck Depression Inventory; EDE-Q = Eating Disorder Examination Questionnaire-6; STAI = State-Trait Anxiety Inventory

## Discussion

This study aimed to examine the effect of short-term fasting on tasks measuring four components of impulsivity. Results showed that, contrary to expectations, participants took longer and opened more boxes in the Fixed Win (FW) condition of the Information Sampling Task (IST), a measure of reflection impulsivity, in the fasted compared to the satiated state. Additionally, short-term fasting was associated with more commission errors during the Affective Shifting Task (AST), indicative of a deficit in action inhibition. When fasted, participants made significantly more errors of commission for food compared to household stimuli during shift blocks. There was no difference between fasted and satiated sessions on the impulsive choices made during the Temporal Discounting Task, or in risky decision-making during the Choice x Risk Task.

Participants opened more boxes and made fewer errors in the Fixed Win (FW) condition of the IST when fasted, indicating a decrease in reflection impulsivity. However, there were no fasted/satiated differences for the Decreasing Win (DW) condition. This suggests that the two conditions were differentially affected by fasting. During the DW condition participants were told that with every box opened, the number of points to be won decreases, hence there is a cost to opening more boxes. However, during the FW condition participants are told that they can open as many boxes as they wish, with no decrease in winnings. An adaptive strategy would be to open all boxes to guarantee a win. However, participants typically guess before all of the boxes have been opened (33).

The results of the IST were contrary to the hypothesis that short-term fasting would be associated with increased reflection impulsivity. The decreased reflection impulsivity displayed during the fasted session could be due to a number of different factors. Firstly, the ability to flexibly shift attention between decision making (deciding which box colour is in the majority), and the action of box opening could be affected by fasting, causing the ‘repetitive’ box opening during the FW condition. This is unlike the DW condition, in which participants are cued by the decreasing points to shift from opening boxes to make a decision about which colour is in the majority. Set-shifting is the process of changing, or switching, between responding to different tasks, rules, or mental sets(31), and has been extensively studied in EDs, (31). Recent research (9) has demonstrated that fasting affects set-shifting, particularly with cue-induced craving (9, 34), and that 18 hours of fasting exacerbates set-shifting difficulties on a rule change task (35). Although this type of short-term fasting in a healthy population is not identical to the patterns of food restriction and chronic or intermittent fasting seen in EDs, it could explain, in part, why participants opened more boxes in the FW condition of the IST when fasted.

Secondly, participants in the fasted session may have become fixated on the detail of opening each box individually and were unable to stand back to see the ‘whole picture’ to make a decision. The term central coherence is used to refer to the ability to combine information into the ‘bigger picture’ rather than focussing only on the finer detail. An impairment in central coherence has been shown in individuals with ED’s (36) and fasted HIs (37). However, an impairment in central coherence may not have occurred in the DW condition as participants may have been cued into making a decision by the decreasing points.

However, it is not possible to determine the contribution of either of these explanations from the current experiment. Therefore the results require further investigation and replication to understand the mechanisms underpinning the effect of decreased reflection impulsivity on the IST.

Results from the current study indicate that short-term fasting did not affect delay aversion. Participants in the fasted condition did not choose to delay the receipt of a monetary reward any less than when satiated. However, participants may have been less susceptible to the fasting manipulation as the hypothetical on-screen choices are viewed as more distant, compared to immediate presentation, and are more objectively assessed (38). The degree to which an individual discounts future rewards has also been described as a trait characteristic (39), and is stable over time (40, 41). Therefore manipulating the state of the individual (fasting) may not influence an established trait discount rate towards monetary rewards.

Participants also showed no difference between fasted and satiated sessions for the different probabilities of winning, different magnitudes of expected losses, and expected gains on the Choice x Risk Task. This indicates that risky decision-making was not influenced by short-term fasting. This finding is in contrast to previous research that found increased risky decision-making for food, water, and money following four hours of food and water deprivation [36]. However, this could be related to differences in the salience of the reward as participants in the current study received points rather than food, water, or money, which may be differentially affected by fasting. Additionally, exploratory analysis of fasted state on risk preferences in Levy and colleagues’ study revealed a small effect (5% change) that appeared to be related to the baseline characteristics of the included sample [36].

Another study demonstrated that risky decision-making decreased when fasted participants were provided with a meal to reach satiation. However, this study involved exclusively male participants [37], whereas, the participants in the current study were all female. Hence, gender differences might account for the inconsistent results, especially when males and females have been shown to respond to fasting differently (42). Furthermore, the effect on risky decision-making in the previous study was only significant immediately after a satiated meal but not one hour later [37].This appears to be in line with the current lack of effect of fasting given that participants in the current study were told to eat normally prior to the satiated session, and were not provided with food during task completion which took between 30 and 60 minutes.

Participants exhibited more errors of commission for food stimuli during the AST when fasting compared to when satiated, indicating a deficit of action inhibition. However, there were no differences in response times between fasted and satiated sessions. The increased number of errors of commission in the fasted condition indicated decreased action inhibition. Higher errors of commission, or decreased action inhibition, in BN compared to HIs have previously been interpreted as indicative of greater impulsivity (21). Participants made significantly more commission errors when fasted during the more difficult shift blocks for food compared to household stimuli. This difference was not present in the non-shift blocks. This result could indicate that participants are less able to control motor impulsivity during a more demanding task, and towards food stimuli when fasted.

Therefore the current findings suggest that short-term fasting may be an important consideration when examining differences in action inhibition between HIs and BN. If individuals with BN undergo periods of short-term fasting, and have a similar response to HIs in the current study, then the increased commission errors in BN could be attributed to fasted state, rather than reflecting an impulsive neurocognitive profile, or trait. It is important to disentangle the contribution of short-term fasting to impulsivity seen in BN so that treatments that focus on reducing impulsivity such as Dialectical Behaviour Therapy can be appropriately informed and targeted.

A limitation of the current experiment is the inability to address whether the differences found between fasted and satiated sessions is due to the primary effect of lowered blood glucose on brain function, or the secondary effect of hunger (induced through fasting) influencing motivation, or fatigue. Previous research indicates that changes in cognition can be independent of blood glucose, and may be mediated by other factors (43), and could be controlled by homeostatic mechanisms not assessed in the current study (44).

Green and colleagues have previously found that although there was a significant difference between self-reported hunger for fasted and satiated sessions, task performance was not affected. This indicates that subjective measures of hunger may not always relate to differences in task performance. The tasks in the current study for which there were non-significant findings may not have sensitive enough to detect subtle differences in performance that could occur as a result of fasting(45). Further research is needed in order to examine the role of subjective hunger on cognition and to separate out the influence of primary and secondary effects of fasting on cognitive performance.

Furthermore, the fasting manipulation might not have increased the value of a monetary reward, but instead increased the value of a food reward. Previous studies have demonstrated that nicotine deprivation can lead to a steeper discounting rate for cigarettes, but not monetary rewards (46). This demonstrates that state manipulations can have differential effects on the impulsive choices made in response to different rewards. The present findings are therefore only applicable to monetary rewards, and future studies should investigate food rewards using this paradigm. This could also account for the non-significant findings during the delay aversion and risky-decision making task, which used monetary values as rewards. However, the present results show that general delay aversion towards money did not differ as a function of fasting. Including food stimuli during the temporal discounting task could make the results difficult to interpret. It might be hard to separate impulsiveness towards food items as a result of fasting from the increased value of food items caused by food deprivation.

It is clear that further studies need to be conducted in order to better understand the effect of short-term fasting in healthy participants. Research should continue to investigate the most appropriate design in which to examine the role of short-term fasting on cognitive performance. In the meantime, caution should be used when interpreting findings from ED participants, particularly BN, as indicative of trait differences in cognitive performance due to the influence of fasted state on these measures.

